# GSK3 inhibition ameliorates the abnormal contractility of Newfoundland ACM patient iPSC-cardiomyocytes

**DOI:** 10.1101/2024.02.06.579108

**Authors:** Rebecca Noort, Wesam Salman, Camila Fuchs, Ursula Braun, David Pace, Kathleen Hodgkinson, Jessica Esseltine

## Abstract

Arrhythmogenic Cardiomyopathy (ACM) is clinically characterized by ventricular arrhythmias causing sudden cardiac death and fibrofatty replacement of the myocardium leading to heart failure. One form of ACM is highly prevalent in the Canadian Province of Newfoundland and Labrador (NL) and has earned the moniker, The “Newfoundland Curse”. This ACM in NL patients is often caused by a fully penetrant heterozygous missense pathogenic variant in the *TMEM43* gene (*TMEM43* c.1073C>T; TMEM43 p.S358L). Although the causative variant has been identified, little is known about the function of the TMEM43 protein in cardiomyocytes, how the TMEM43 p.S358L mutation contributes to the development of arrhythmias, or why the disease is more severe in males than females. To explore the role of TMEM43 in cardiomyocyte function, we generated induced pluripotent stem cells (iPSCs) from 2 severely affected male Newfoundland ACM (TMEM43 p.S358L) patients. CRISPR-Cas9 was used to genetically “repair” the heterozygous *TMEM43* variant in ACM patient iPSCs or for *TMEM43* gene knockout. ACM patient iPSC-cardiomyocytes with the TMEM43 p.S358L variant display pro-arrhythmogenic phenotypes *in vitro* with significantly elevated contraction rates and altered calcium handling, although no obvious gross abnormalities were observed across several major intracellular organelles. GSK3 inhibition significantly increased protein expression of β-catenin as well as Lamin A/C and ameliorated the pro-arrhythmic tendencies of ACM patient iPSC-CMs.

## Introduction

Arrhythmogenic Cardiomyopathy (ACM) is a category of related heritable heart diseases featuring both ventricular arrhythmias causing sudden cardiac death (SCD) and cardiomyopathy leading to heart failure^1^. About 60% of ACM cases have known genetic links across at least 13 different genes^2, 3^. ACM is often associated with the dysfunction of the desmosomes, which provide strength to the cardiac muscle. When the desmosomes break down, the myocardium cannot withstand the pressures exerted during beating, causing cardiomyocytes to break apart and die^1^. Cardiomyocyte electrical coupling via gap junctions is subsequently compromised when non-excitable fibrous tissue invades the damaged myocardium leading to ventricular arrhythmia. Other ACM pathogenic variants involve various calcium handling proteins as well as nuclear envelope proteins and linker of nucleoskeleton and cytoskeleton (LINC) complex^4^. In 2008, researchers at Memorial University of Newfoundland found an autosomal dominant mutation in the *TMEM43* gene (*c.1073C>T*, p.S358L) within a cluster of patients suffering from ACM^5, 6^. The TMEM43 p.S358L variant was subsequently reported in the UK^7^, Germany^8^, Denmark^9^ and Japan^10^. This pathogenic gene variant is fully penetrant, thus every individual who inherits the disease-causing allele will develop at least one clinical feature of ACM^6, 11^. TMEM43 p.S358L-linked ACM is an autosomal dominant disorder, therefore children born into these families have a 50% risk of inheriting the disease-causing *TMEM43* allele. TMEM43 p.S358L ACM is significantly more severe in males than females: 50% of untreated males die by 40 years, 80% by 50 years (5% and 20% respectively for females)^1, 11^. Along with sex, the greatest risk factor for severe disease presentation is exercise^12–14^. This risk has lead to NL pediatric cardiology supporting early genetic testing and restriction of high intensity sports for children with the TMEM43 p.S358L pathogenic variant^14^. Primary ACM treatment for TMEM43 p.S358L ACM individuals is the Implantable Cardioverter Defibrillator (ICD), the use of which has added an average 18 years of life to TMEM43 ACM patients compared to their untreated ancestors^5^. A large cohort of individuals with the TMEM43 p.S358L variant and their siblings and relatives born at an *a priori* 50% risk are ascertained in NL, with extensive clinical data over several generations.

Although TMEM43 p.S358L is a cause of ACM, we know very little regarding the function of TMEM43 protein or how *TMEM43* pathogenic variant compromises cardiomyocyte function. Here we generated and characterised induced pluripotent stem cell-cardiomyocytes (iPSC-CMs) from 2 severely affected male Newfoundland ACM patients with the autosomal dominant TMEM43 p.S358L mutation. iPSCs are ideal for disease modeling as they are of human origin, are patient-specific, amenable to genetic manipulation, and can differentiate into various terminal cell types. Unaffected familial controls together with isogenic “repair” iPSCs revealed that ACM iPSC-CMs exhibit profound pro-arrhythmic tendencies and rapid spontaneous contractions that can be ameliorated with GSK3 inhibition. The patient-oriented nature of this research will help inform some of the most fundamental mechanisms of cardiomyocyte dysfunction in patients with TMEM43 p.S358L pathogenic variant.

## Material and Methods

### Human Ethics

These studies are in accordance with the Declaration of Helsinki and were approved by the Newfoundland and Labrador Human Research Ethics Board (HREB #2018.210 & 2020.301). Written consent was secured prior to enrolling subjects in the study. All approvals are reviewed and renewed annually.

### ACM Patient Dermal Biopsy and Primary Fibroblast Isolation

The patient samples used in this study included two severely affected male ACM patients (ACM1 and ACM2) from NL as well as one unaffected brother (sibling of ACM1). Primary fibroblasts were isolated via outgrowth from dermal punch biopsies. Primary fibroblasts were cultured on 0.1% gelatin coated dishes in DMEM supplemented with FBS (20% v/v). Cells were maintained in an incubator at 37°C and 5% CO_2_. The cells were passaged with Trypsin-EDTA once they reached confluency. Passage 2-3 primary dermal fibroblasts were used for iPSC reprogramming.

### Human Induced Pluripotent Stem Cell Reprogramming & Characterization

Dermal fibroblasts were reprogrammed using a ReproRNA™-OKSGM Kit (Cat # 05903, STEMCELL Technologies, Vancouver, CAN) according to the manufacturer’s instructions. Resulting iPSC colonies were picked and transitioned onto Geltrex-coated plates with Essential 8 medium. At passage 8, iPSC lines were tested for mycoplasma contamination (Mycoplasma PCR Detection Kit; Cat # G238, abm, Richmond, CAN), normal copy number at mutation hotspots (hPSC Genetic Analysis Kit; Cat # 07550, STEMCELL Technologies), and >90% expression undifferentiated state markers (TRA-

1-60, SSEA4, OCT4; see antibodies in Table 1). *TMEM43* variant status in the iPSCs was confirmed relative to the starting fibroblast population by Sanger Sequencing a PCR amplicon flanking the c.1073 mutation site. Only resulting iPSC lines that passed all quality checks were used for experiments.

**Table 1.** Antibodies and Dyes used in Western Blot, Immunofluorescence, & Flow Cytometry

| Marker | Supplier | Catalog # | Host Species | IF | Western Blot | FC |
| --- | --- | --- | --- | --- | --- | --- |
| 6X-His Tag | ThermoFisher Scientific | MA5-33032 | rabbit |  | 1:1000 |  |
| AKT (pan) | Cell Signaling Technology, (Danvers, USA) | 4691 | rabbit |  | 1:1000 |  |
| Calnexin | Cell Signaling Technology | 2679 | rabbit |  | 1:1000 |  |
| Calnexin | ThermoFisher Scientific | MA3-027 | mouse |  | 1:1000 |  |
| Connexin 43 | MilliporeSigma (Burlington, USA) | C6219 | rabbit | 1:1000 | 1:5000 |  |
| cTNT (cardiac troponin T) | Abcam (Cambridge, GBR) | ab8295 | mouse | 1:800 |  | 0.1 uL/test |
| Emerin | Abcam | ab204987 | mouse |  | 1:1000 |  |
| Emerin | Cell Signaling Technology | 30853 | rabbit |  | 1:1000 |  |
| FLAG Tag | MilliporeSigma | F1804 | mouse |  | 1:1000 |  |
| FLAG Tag | Cell Signaling Technology | 14793 | rabbit |  | 1:1000 |  |
| GAPDH | MilliporeSigma | MAB374 | mouse |  | 1:5000 |  |
| Lamin A/C | Abcam | ab238303 | mouse | 1:1000 | 1:1000 |  |
| MLC2V | Abcam | ab79935 | rabbit | 1:500 |  |  |
| N-Cadherin | BD Biosciences (Franklin Lakes, USA) | 610921 | mouse | 1:500 | 1:2000 |  |
| OCT4 BrilliantViolet™421 | BioLegend (San Diego, USA) | 653712 | mouse |  |  | 0.5 uL/test |
| Phospho-AKT (Ser473) | Cell Signaling Technology | 4060 | rabbit |  | 1:1000 |  |
| Phospho-AKT (Thr308) | Cell Signaling Technology | 13038 | rabbit |  | 1:1000 |  |
| Phospho-c-Raf (Ser259) | Cell Signaling Technology | 9421 | rabbit |  | 1:1000 |  |
| Phospho-GSK-3β (Ser9) | Cell Signaling Technology | 5558 | rabbit |  | 1:1000 |  |
| Phospho-PDK1 (Ser241) | Cell Signaling Technology | 3438 | rabbit |  | 1:1000 |  |
| Phospho-PTEN (Ser380) | Cell Signaling Technology | 9551 | rabbit |  | 1:1000 |  |
| Plakoglobin (gamma catenin) | Abcam | ab231304 | mouse | 1:1000 |  |  |
| SERCA2 / ATP2A2 | Cell Signaling Technology | 9580 | rabbit |  | 1:1000 |  |
| SSEA4 AF647 | BioLegend | 330408 | mouse |  |  | 0.2 uL/test |
| TMEM43 | Abcam | ab184164 | rabbit |  | 1:2000 |  |
| TRA-1-60-R | BioLegend | 330614 | mouse |  |  | 2 uL/test |
| Goat-anti-mouse IgG, HRP-linked | ThermoFisher Scientific | 31430 | Goat |  | 1:5000 |  |
| Goat-anti-rabbit IgG, HRP-linked | Cell Signaling Technology | 7074 | Goat |  | 1:1000 |  |
| donkey-anti-mouse AF555 | ThermoFisher Scientific | A31570 | Donkey | 1:1000 |  |  |
| donkey-anti-rabbit AF488 | ThermoFisher Scientific | A21206 | Donkey | 1:1000 |  |  |
| goat-anti-mouse AF647 | ThermoFisher Scientific | A32728 | Goat | 1:1000 | 1:5000 | 1:3000 |
| goat-anti-rabbit AF488 | ThermoFisher Scientific | A78953 | Goat |  | 1:5000 |  |
| Hoechst 33342 | FisherScientific | H3570 | Dye | 1:1000 |  |  |
| Phalloidin-647 | Cell Signaling Technology | 8940 | Dye | 1:20 |  |  |
| Zombie-Aqua fixable viability dye | BioLegend | 423101 | Dye |  |  | 1:1000 |
| Zombie-NIR fixable viability dye | Biolegend | 423105 | Dye |  |  | 1:1000 |
| Mouse IgG1 isotype control | Cell Signaling Technology | 5415 | mouse |  |  |  |

### Human Induced Pluripotent Stem Cell Culture

iPSCs were maintained in either Essential 8 medium (E8) (Cat # A1517001, ThermoFisher Scientific) or mTeSR™-Plus (Cat # 100-0276, STEMCELL Technologies) in 6-well plates pre-coated with Geltrex (Cat # A1413302, ThermoFisher Scientific). Cells received daily media changes and were passaged every 4-5 days as aggregates using 0.5 mM EDTA in DPBS (Cat # AM9260G, ThermoFisher Scientific). Cell aggregates were gently scraped off the bottom of the well and re-seeded onto a new Geltrex-coated dish.

### TMEM43 p.358L CRISPR-Cas9 Isogenic Repair

iPSCs from patient ACM1 harbouring the pathogenic *TMEM43* c.1073T (p.358L) allele were corrected to *TMEM43* c.1073C (p.358S) to create isogenic repair iPSCs. CRISPR-Cas9 gene editing was performed according to STEMCELL Technologies’ *Genome Editing of Human Pluripotent Stem Cells Using the ArciTect™ CRISPR-Cas9 System* Technical Document # 27084 Version 3.0.1 FEB 2022 with slight modification in conjunction with the Neon™ Electroporation System (Cat # MPK5000, ThermoFisher Scientific).

A gRNA proximal to the *TMEM43* c.1073T mutation site was identified with the Sanger Institute CRISPR finder (CRISPR ID 950613618 (5’- CACGGTCAGCAGGGTCAGCG-3′)) and was modified by changing the 3’ guanine to an adenine to make the gRNA specific to the pathogenic *TMEM43* allele. Analysis of the gRNA with IDT’s CRISPR-Cas9 guide RNA design checker reveals no probable off-target effects. A 108 nt ssODN (single stranded oligodeoxynucleotide) with symmetrical homology arms was created using IDT’s Alt-R™ CRISPR HDR Designing Tool and the positive strand polarity was selected based on principles described^15^. The ssODN contains three altered nucleotides to (A) correct *TMEM43* c.1073T to c.1073C, (B) destroy the PAM sequence after cutting, and (C) ablate an MlsI digest site.

Low-passage ACM iPSCs with normal chromosome copy number were subjected to CRISPR-Cas9 gene editing by electroporation with an RNP (ribonucleoprotein) complex and the ssODN along with a GFP-containing plasmid to facilitate identification of successfully transfected cells. The electroporation reaction consists of 1.248 µM Cas9, 4 µM sgRNA, 3.33 µM ssODN, 0.033 µg/µL pEGFP-N1 plasmid, and 66,000 single cell iPSCs/µL. The mixture was loaded into a Neon™ 10 µL pipet tip, electroporated with 1 pulse at 1200V for 30 ms, plated into a single well of a Geltrex-coated 24 well plate containing mTeSR™ Plus + CloneR™2 (Cat # 100-0691, STEMCELL Technologies) + 0.1% (v/v) Alt-R(T) HDR Enhancer V2 reagent, and incubated overnight at 37°C at 5% CO_2_.

The following day (14-20 hours after electroporation), the cells were singularized using Accutase and GFP+ cells were isolated by FACS. Sorted cells were plated at clonal density (5-80 cells/cm^2^) onto Geltrex-coated plates containing mTeSR™ Plus and CloneR™2 for three days followed by mTeSR™ Plus thereafter. Well-isolated iPSC colonies were picked and assayed for the desired gene edit by MlsI restriction digest assay and by Sanger sequencing (The Center for Applied Genomics, SickKids Hospital, Toronto, Canada). Resulting isogenic repair iPSCs that passed all quality checks as described above (see Human Induced Pluripotent Stem Cell Reprogramming & Characterization) were used for subsequent experiments.

### Cardiomyocyte Differentiation & Replating

iPSCs were differentiated into contracting ventricular cardiomyocytes using a commercially available kit (STEMdiff™ Ventricular Cardiomyocyte Differentiation kit, Cat # 05010, STEMCELL Technologies). iPSCs were passaged into single cells using Accutase (Cat # A1110501, ThermoFisher Scientific) at a cell density of 2.0 - 2.5 x 10^5^ cells/cm^2^ into the desired vessel. Once the cells reached >95% confluency, cardiomyocyte differentiation was initiated as per the manufacturer’s instructions. Beating cardiomyocytes were fed every other day with STEMdiff^TM^ Maintenance Medium (Cat # 05020, STEMCELL Technologies) until analysis on day 40 or replating.

Contracting cardiomyocytes were singularized for downstream applications like flow cytometry, calcium imaging, fluorescence microscopy, and mitochondrial stress assays. Cells were incubated in a solution of 2% collagenase type II (Cat # LS004174, Worthington Biochem, Lakewood, USA), prepared in Hanks Balanced Salt Solution (HBSS; Cat # 14025092, ThermoFisher Scientific) at 37°C for 20 – 30 minutes to detach the cardiomyocytes from the growth surface. The collagenase solution was removed and the cells were singularized using 0.25% trypsin/ 0.5 mM EDTA for 5-8 minutes at 37°C followed by gentle trituration. Cardiomyocytes were pelleted to remove the trypsin and replated at a density of 1.0 - 2.0 x 10^5^ cells/cm^2^ onto Geltrex-coated dishes containing STEMdiff^TM^ Maintenance Medium + 10 uM Y-27632. The day after replating, medium was changed to STEMdiff^TM^ Maintenance Medium without Y-27632 and cardiomyocytes were permitted to re-establish contractions (2-4 days) before use in downstream assays. Generally, iPSC-cardiomyocytes were used for experimentation on protocol day 40, with approximately 4 weeks of contraction *in vitro*.

### SDS-PAGE & Western Blot

Cells were lysed in ice-cold lysis buffer supplemented with protease and phosphatase inhibitors (150 mM NaCl; 50 mM Tris pH 8.0; 1.0% Triton X-100; 2 µg/mL Leupeptin; 2 µg/mL Aprotinin; 10 mM NaF; 1 mM Na_3_VO_4_; 0.02% NaN_3_). Soluble protein concentration was determined using the Pierce BCA Protein Assay Kit (Cat # PI23225, ThermoFisher Scientific) as per the manufacturer’s instructions. Proteins were separated via sodium dodecyl sulfate-polyacrylamide gel electrophoresis (SDS-PAGE) on 7.5% polyacrylamide gels or 4-20% gradient gels (Cat # 4561094 Bio-Rad, Hercules, USA). The gel was then transferred to a nitrocellulose membrane with a pore size of 0.45 µm (Cat # 1620115, Bio-Rad). The nitrocellulose membrane was incubated in primary antibody diluted in blocking buffer (3% w/v milk in pH 7.6 Tris Buffered Saline with TWEEN® 20 (TBST)): 15.23 mM Tris HCl, 4.62 mM Tris base, 150 mM NaCl, and 0.1% TWEEN® 20. Primary and secondary antibodies used for Western blot were prepared in block buffer as indicated in Table 1. Where used, HRP-conjugated secondary antibodies were visualized using Clarity™ Western ECL Substrate (Cat # 1705061, Bio-Rad) and blot images captured using the ChemiDoc™ MP Imaging System (Cat # 12003154 Bio-Rad). Immunoblots were analyzed by densitometry using FIJI software (ImageJ 1.52p). Every immunoblot was normalized to the house-keeping protein glyceraldehyde 3- phosphate dehydrogenase (GAPDH) when performing densitometric analysis.

### Confocal Microscopy

Cells cultured on #1.5 glass coverslips or Nunc™ Lab-Tek™II chamber slides (Cat # 154453, ThermoFisher Scientific) were washed with DPBS and fixed in 10% normal buffered formalin (Cat # HT501128, MilliporeSigma) for 10 minutes at room temperature. Fixed cells were permeabilized in PBS-T (DPBS + 0.1% TWEEN®20) for 20 minutes followed by 0.1% Triton-X-100 in DPBS for 10 minutes. Primary antibodies were diluted in PBS-T + 5% BSA (Cat # 800-095-EG, Wisent) + 0.05% NaN_3_ and incubated overnight at 4°C. Secondary antibodies and dyes (prepared in PBS-T were incubated at room temperature for 2 hours. Antibodies and stains used for immunofluorescence are indicated in Table 1. The slides were mounted using mowial®488 reagent with 1,4- diazabicyclo[2.2.2]octane (DABCO) antifade compound as described by Cold Spring Harbor Protocols^16^.

Immunofluorescence confocal images were taken on a Zeiss LSM 900 with Airyscan 2 fitted with a 63X/1.4NA oil immersion lens and the following lasers: 405 nm, 488 nm, 561 nm, 640 nm. All images taken were analyzed and processed using FIJI software.

### Electron Microscopy

Contracting cardiomyocytes were rinsed with DPBS and scraped as a single sheet from the culture surface and fixed in Karnovsky’s fixative overnight. Fixed samples were processed for transmission electron microscopy according to standard methods. Images were captured on a Tecnai Spirit Transmission electron microscope with a 4-megapixel AMG digital camera.

### Flow Cytometry

Single cell suspensions were prepared for flow cytometry and stained with Zombie fixable viability dyes and antibodies as shown in Table 1. Samples were analyzed on a CytoFLEX flow cytometer (Beckman Coulter) equipped with 405 nm, 488 nm, and 638 nm lasers. Analysis was performed with FlowJo™ v.10.3 Software. UltraComp eBeads™ Plus Compensation Beads (Cat # 01-3333-41, ThermoFisher Scientific) labeled with fluorescent-tagged antibodies were used for compensation. Gate placement was determined using fluorescence minus one (FMO) or secondary antibody only controls as appropriate.

### Live Cell Calcium Imaging

Contracting iPSC-CMs cultured on high end microscopy µ-slides (Cat # 80806, Ibidi, DEU) were incubated with 10 µM Fluo-4 AM dye (Cat # F14201, ThermoFisher Scientific) in cardiomyocyte maintenance medium for 15 minutes at 37°C and 5% CO_2_. After incubation, iPSC-CMs were washed with HBSS (Cat # 14025092, ThermoFisher Scientific) and fresh cardiomyocyte maintenance medium was then added to wells. Calcium transients were captured on a Zeiss LSM 900 with Airyscan 2 with a 20X/0.8NA objective and 488 nm laser. The cells were housed in a humidified 37°C chamber with 5% CO_2_ throughout image acquisition.

### Drug Treatments

Contracting iPSC-CMs (at protocol day 11 – 20) were treated with STEMdiff™ Maintenance Medium supplemented with 2 µM of the GSK3 inhibitor CHIR99021 (Cat # 13122, Cayman Chemicals, Ann Arbor, USA) for ∼20 consecutive days until analysis on day 40.

### Calcium Imaging Analysis

Images taken over a 20-30 second time period were analyzed by FIJI with the “plot profile” to generate a list of xy coordinates for Fluo-4 AM intensity over time. These values were processed using Origin® 2023 software (OriginLab Corporation, Northhampton, USA) with the calcium transient app to determine F/F0 and other parameters describing the shape of each calcium transient. Data were plotted using GraphPad Prism version 9.4.1 (GraphPad Software Inc, San Diego, USA).

### Seahorse Mitochondrial Stress Assay

Mitochondrial function of iPSC-CMs with or without GSK3 inhibition was evaluated with the Seahorse XF Cell Mito Stress Test Kit (Cat # 103015-100, Agilent) in conjunction with Seahorse XFe24 FluxPak mini (Cat # 102342-100, Agilent). Contracting iPSC-CMs on protocol day ∼32 were singularized as described above and seeded at 2.0 x 10^5^ viable cells per well of a Geltrex-coated Agilent Seahorse XF24 microplate (Cat # 100777-004) into plating medium (STEMdiff Maintenance Medium + 10 µM Y-27632). For CHIR99021- treated iPSC-CMs, 2 uM CHIR99021 was added to the plating medium. iPSC-CMs were cultured on the XF24 microplate for an additional 8 days in STEMdiff Maintenance Medium M +/- CHIR99021 until the cells reached day 40. The mitochondrial stress assay was performed according to the instructions in the Agilent Seahorse XF Mito Stress Test Kit User Guide (manual 103016-400, second edition, May 2019) using XF DMEM medium (Cat # 103575-100) supplemented with 10 mM glucose, 1 mM pyruvate, and 2 mM glutamine. Measurements were performed on an Agilent Seahorse XFe24 Analyzer.

### Statistical Analysis

All statistical analyses were performed using GraphPad Prism version 9.4.1 (GraphPad Software Inc, San Diego, USA). Bar graph data are presented as mean ± standard error values and violin plots show the median and quartile positions. When comparing two data sets, unpaired t-test was used to analyze data and determine significance. For data sets of three or more, one-way analysis of variance (ANOVA) with Tukey’s post hoc multiple comparisons test was used to determine statistical significance. A p-value of less than 0.05 was considered significant in this study where not significant (ns), p>0.05, * p≤0.05, ** p≤0.01, *** p≤0.001, and **** p≤0.0001.

## Results

### ACM patient iPSC-cardiomyocytes exhibit pro-arrhythmic tendencies

In this study we reprogrammed iPSCs from 2 male NL ACM patients with the TMEM43 p.S358L heterozygous pathogenic variant (ACM1 and ACM2). These individuals were selected based on a severe disease presentation: ICD discharge for sustained VT ≥ 240 bpm under age 35 years and a maintained history of appropriate discharges and progression of disease ultimately requiring heart transplant before the age of 40 (Table 2). We also reprogrammed iPSCs from one unaffected brother (sibling of ACM1). CRISPR-Cas9 genetic engineering was used to create isogenic control cells by “repairing” the TMEM43 p.S358L mutation in ACM1 patient iPSCs (Figure 1 A-C).

**Figure 1.**
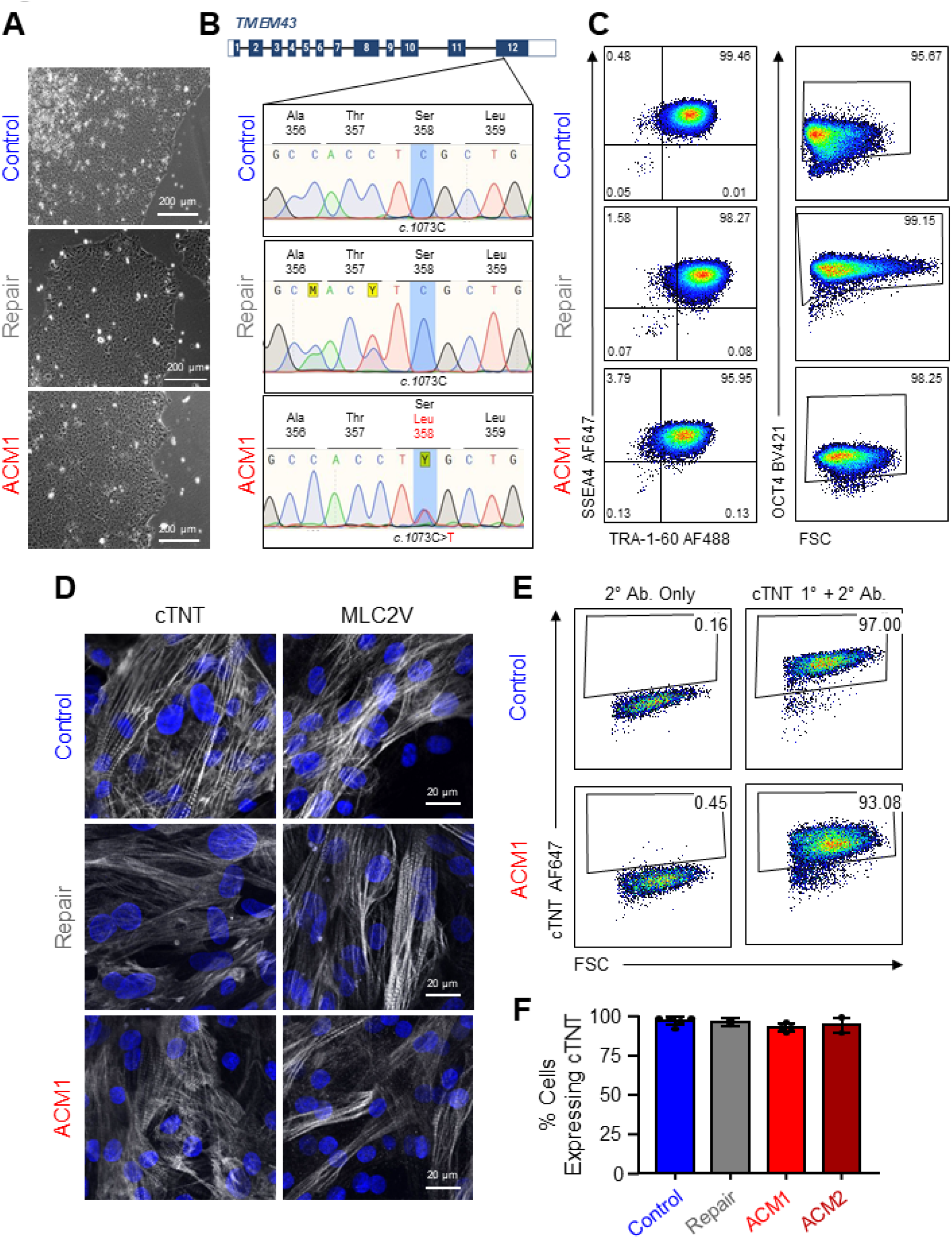
ACM iPSC reprogramming, CRISPR gene editing, and cardiac differentiation. **(A)** Representative phase contrast micrograph demonstrating typical morphology of iPSCs reprogrammed from control and ACM patient donors. **(B)** Representative flow cytometric plot showing high expression of key pluripotency markers in our iPSCs. **(C)** Gene schematic and Sanger Sequencing result of unaffected control, ACM patient and CRISPR-Cas9 isogenic repair iPSCs. In addition to repairing the C>T transition, the repair template included silent mutations to destroy the CRISPR PAM site (eliminate repeated Cas9 cutting) and delete a restriction enzyme cut site (facilitate screening). **(D)** Representative immunofluorescence confocal micrographs demonstrating cTNT and MLC2V expression in our differentiated iPSC-cardiomyocytes. **(E, F)** Representative flow cytometric plot and associated quantification of the proportion of cTNT expressing cells after iPSC-cardiomyocyte differentiation.

**Table 2.**
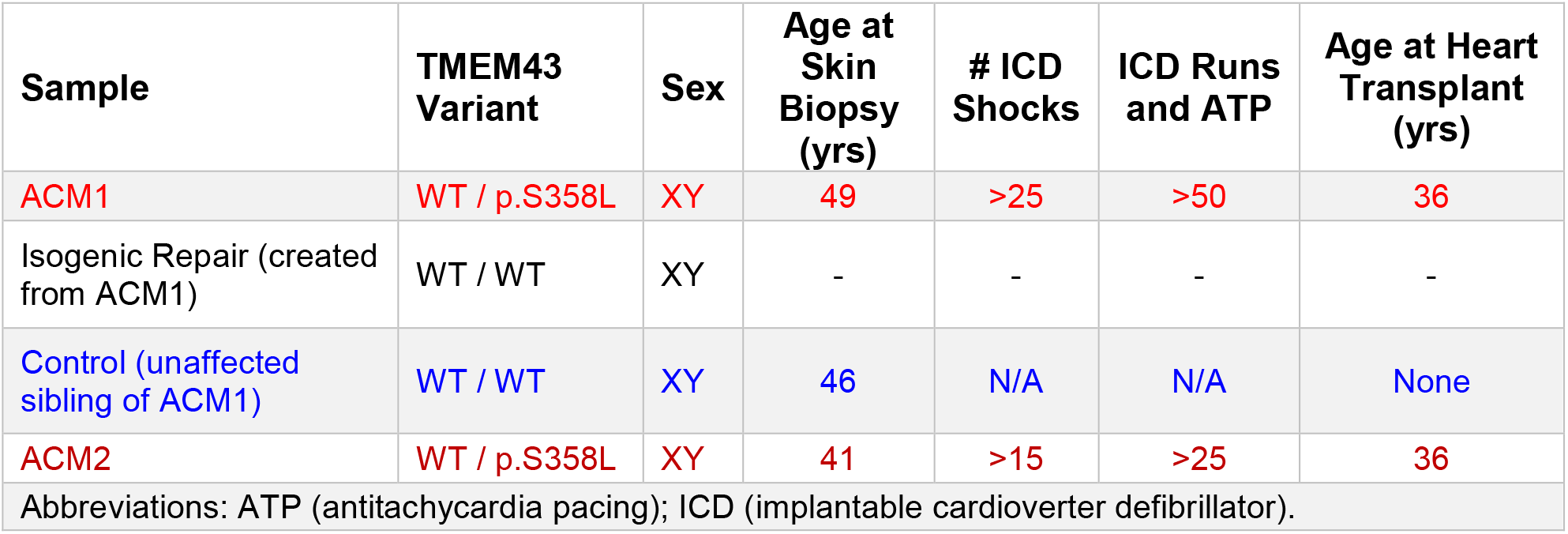
Patient-derived iPSC samples included in the study

All iPSC lines successfully differentiated into cardiomyocytes and routine testing was used to confirm high proportions of differentiated cells expressing the cardiomyocyte marker cardiac troponin T (cTNT) and the ventricular myocyte marker myosin light chain 2V (MLC2V) (Figure 1 D-F). It became immediately apparent that both ACM patient-derived iPSC-CM lines exhibited significantly higher spontaneous contraction frequencies under basal conditions (Figure 2 A-C, Supplemental videos). Indeed, our unaffected control iPSC-CMs contracted an average of 16.25 +/- 0.6226 times per minute while the ACM iPSCs contracted an average of 39.07 +/- 2.539 (patient ACM1) or 38.09 +/- 2.161 times per minute (patient ACM2). This increased contraction rate resulted in a significant decrease of nearly all of the calcium transient parameters tested, including time to peak, decay time, transient duration and time between maxima (Figure 2 C). Moreover, blinded scoring showed a dramatic increase in the number of pro-arrhythmic events from ACM patient iPSC-CM calcium traces compared to control or isogenic repair iPSC-CMs (Figure 2 D). Blinded classification of the arrhythmic traces (Figure 2 E) reveals the majority present as alternans-like, followed by early afterdepolarization (EAD)-like, and non-uniform inter-event intervals (non-uniform spacing between single, adjacent peaks).

**Figure 2.**
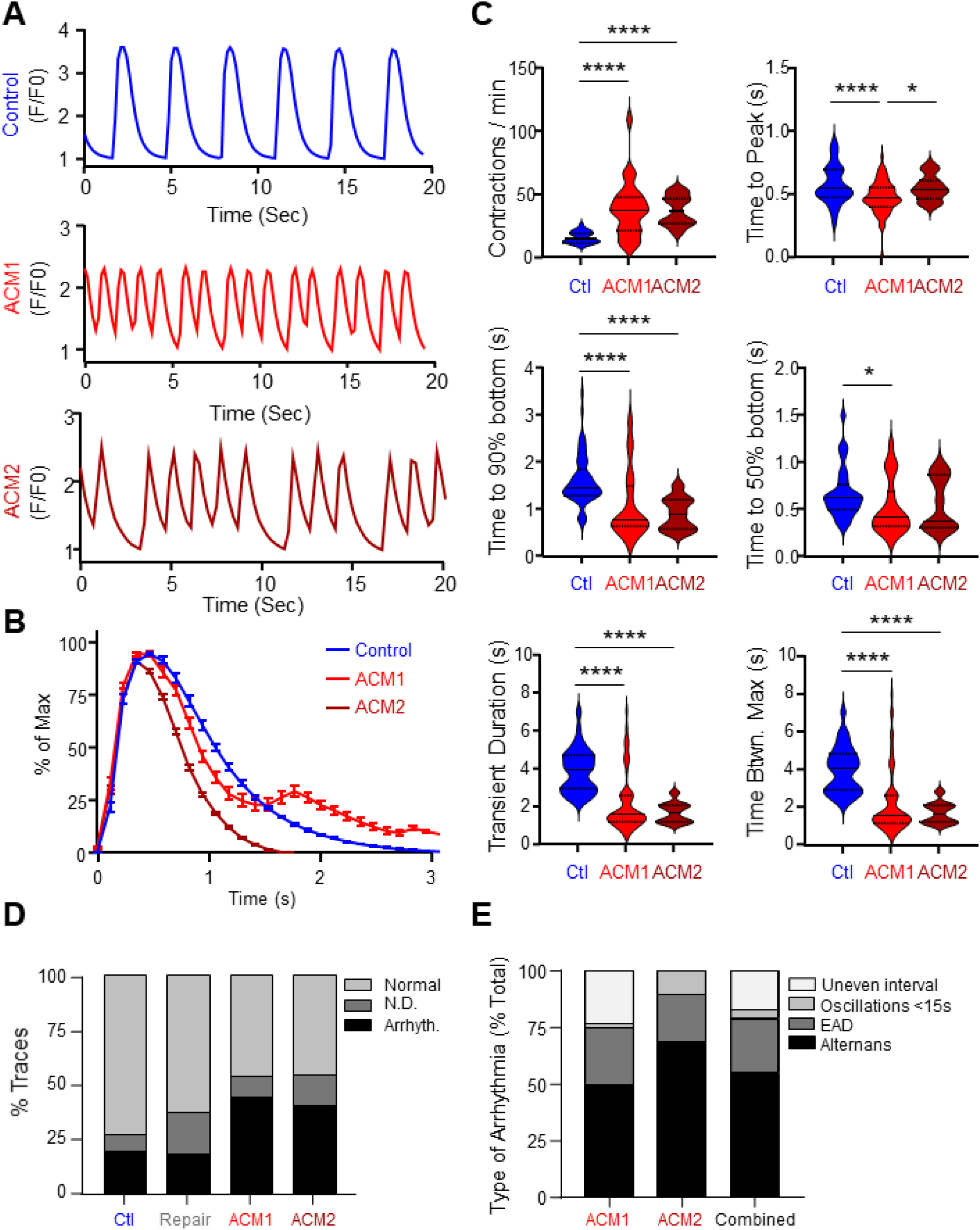
ACM patient iPSC-cardiomyocytes exhibit pro-arrhythmic tendencies. **(A)** Representative Fluo4 traces demonstrating calcium handling in unstimulated iPSC-CMs after 4 weeks of contraction in culture. **(B)** Average calcium peak shape as determined by overlaying all peaks. **(C)** Quantified parameters associated with calcium traces. **(D)** Blinded scoring of calcium image tracing. Traces were scored either as “normal”, pro-arrhythmic” (Arrhyth.) or “not determined” (N.D). **(E)** Blinded scoring of type of arrhythmic event. Pro-arrhythmic traces were scored as either alternans, early afterdepolarizations (EAD), oscillations lasting less than 15 seconds or uneven event interval. Data representative of 3-8 independent experiments, comprising 29-63 individual calcium traces for each cell line. n.s, not significant; *, p < 0.05; **, p < 0.01; ***, p < 0.001; ****, p < 0.0001 as determined using one-way ANOVA with Tukey’s post-hoc multiple comparisons test.

### ACM patient iPSC-CMs exhibit typical cellular organelle morphology

Several other ACM-causing mutations have been reported to impact various intracellular organelles including gap junctions, desmosomes, nuclear envelop proteins and others^2, 3^. Thus, we sought to determine whether any of these organelles were obviously perturbed in our ACM patient iPSC-CMs. Immunofluorescence confocal imaging and electron microscopy both revealed little apparent differences in nuclear envelope, gap junctions, desmosomes, sarcomeres, Z-discs, mitochondria structure, or protein localization between the unaffected control and ACM patient iPSC-CMs (Figure 3 A-C). Finally, flow cytometric analysis of the apoptosis marker cleaved caspase 3 (CC3) was used to determine whether ACM patient iPSC-cardiomyocytes exhibited increased apoptosis rates (Figure 3 D). We observed similarly low rates of CC3-positive cells across all of the cell lines examined (< 1%), suggesting that our ACM patient iPSC-cardiomyocytes do not exhibit increased apoptosis rates *in vitro*.

**Figure 3.**
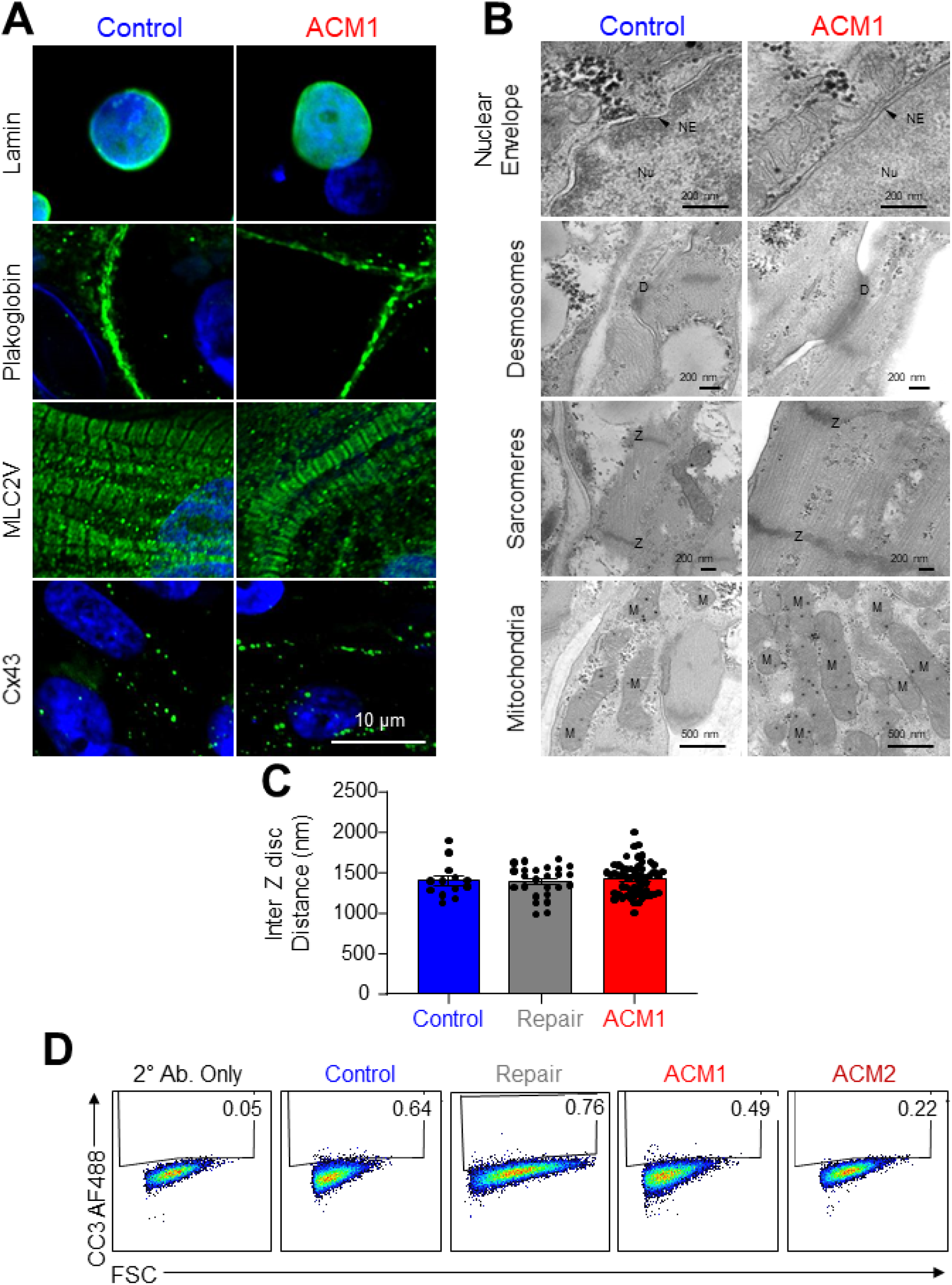
ACM patient iPSC-cardiomyocytes exhibit typical cellular organelle morphology. **(A)** Representative immunofluorescence confocal micrographs and **(B)** representative transmission electron micrographs of various intracellular organelle markers (green) in unaffected control and ACM patient iPSC-cardiomyocytes. Nuclei (Hoechst, blue). Scale bars as indicated. ER, endoplasmic reticulum; NE, nuclear envelope; Nu, nucleus; D, desmosome; Z, Z-disc; M, mitochondria. **(C)** Quantification of the distance between Z-discs in control and ACM patient iPSC-cardiomyocytes. Measurements were made using electron microscopy images. **(D)** Representative flow cytometric analysis plots for cleaved capsase 3 (CC3; marker of apoptosis) in unaffected control and ACM patient iPSC-cardiomyocytes.

### ACM iPSC-CM pro-arrhythmic tendencies can be ameliorated with GSK3 inhibition

Altered signal transduction and/or metabolism has also been proposed as a mechanism of ACM associated with TMEM43 p.S358L^17^. However, we found no difference in key signaling proteins within the AKT signaling axis between ACM patient and unaffected control iPSC-CMs, including phospho-AKT, phospho-GSK3β and others (Figure 4 A-B). Although we observed no changes in phospho-GSK3β protein expression in our ACM iPSC-CMs, others have shown that inhibition of GSK3β can reduce cardiac conduction, ameliorate cardiac dysfunction, and reduce contraction abnormalities^17–19^ Indeed, treatment with the GSK3 inhibitor CHIR99021 had a profound effect on ACM1 iPSC-CMs, improving every metric we evaluated including reduced contraction frequency, extended transient duration, and vastly reduced pro-arrhythmic traces in blinded scoring (Figure 5 A-E).

**Figure 4.**
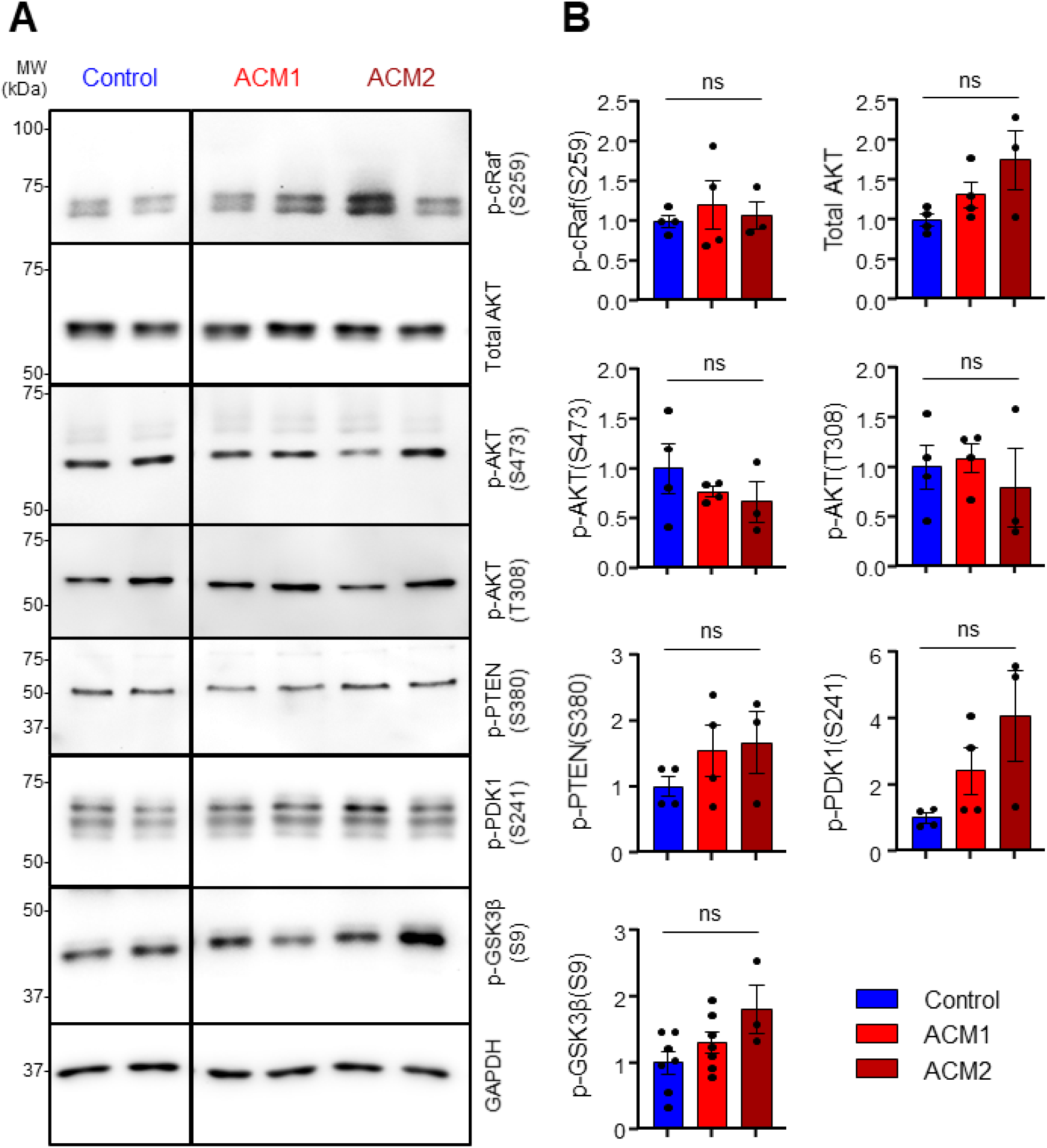
AKT signaling pathway unperturbed in ACM iPSC-cardiomyocytes. **(A)** Representative Western blots and **(B)** densitometric analysis of various signaling molecules in unaffected control and ACM patient iPSC-CMs. Data represent the standard error of the mean of 4 independent experiments.

**Figure 5.**
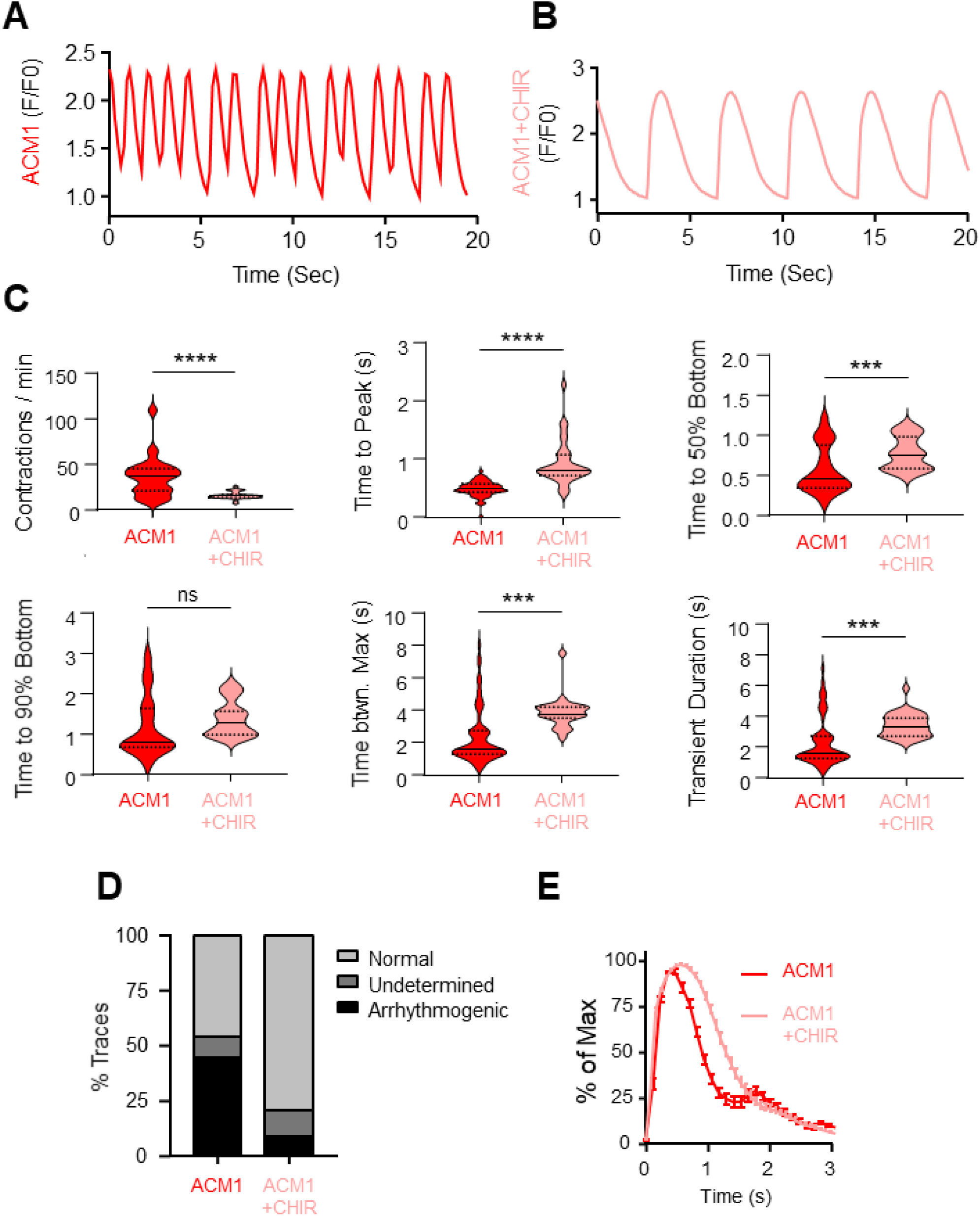
ACM iPSC-cardiomyocyte pro-arrhythmic tendencies can be ameliorated with GSK3 inhibition. (A, **B)** Representative Fluo4 traces demonstrating calcium handling in untreated ACM patient iPSC-CMs or cells treated with 2 µM of the GSK3 inhibitor CHIR99021. **(C)** Quantified parameters associated with calcium traces in CHIR- treated ACM patient iPSC-CMs. **(D)** Blinded scoring of calcium image tracing. Traces were scored either as “normal”, pro-arrhythmic” (Arrhyth.) or “not determined” (N.D). **(E)** Average calcium peak shape of CHIR-treated ACM patient iPSC-CMs. Data representative of 3 independent experiments, comprising 24-34 individual calcium traces for each cell line. n.s., not significant; ***, p < 0.001; ****, p < 0.0001 compared to untreated as determined using one-way ANOVA with Tukey’s post-hoc multiple comparisons test.

Given that GSK3 inhibition decreases spontaneous contraction frequency of our ACM1 iPSC-CMs we wondered if mitochondrial function was impacted by CHIR99021 treatment. To that end, we used a commercial mitochondrial stress test to evaluate control and ACM1 iPSC-CMs with and without GSK3 inhibition (Figure 6). All conditions tested displayed similar profiles for oxygen consumption rate (OCR | Figure 6 A) and extracellular acidification rate (ECAR | Figure 6 B) in response to modulators of mitochondrial respiration. While we did observe a significant difference for non-mitochondrial oxygen consumption between control and ACM1+CHIR iPSC-CMs, all other measured parameters were comparable (Figure 6 C). Since mitochondrial function was largely unaltered, we next investigated whether CHIR99021 treatment influences protein expression levels in ACM patient iPSC-CMs (Figure 7). Western blot analysis showed comparable levels of TMEM43 and of phospho-GSK3β (inactive form) across control and ACM1 iPSC-CMs with or without CHIR99021 treatment. We next examined adherens junction proteins N-cadherin and β-catenin. N-cadherin was not significantly different but β-catenin was significantly upregulated in CHIR99021-treated ACM1 cells relative to untreated iPSC-CMs. Interestingly, we observed a significant increase in lamin A and C expression following CHIR99021 treatment.

**Figure 6.**
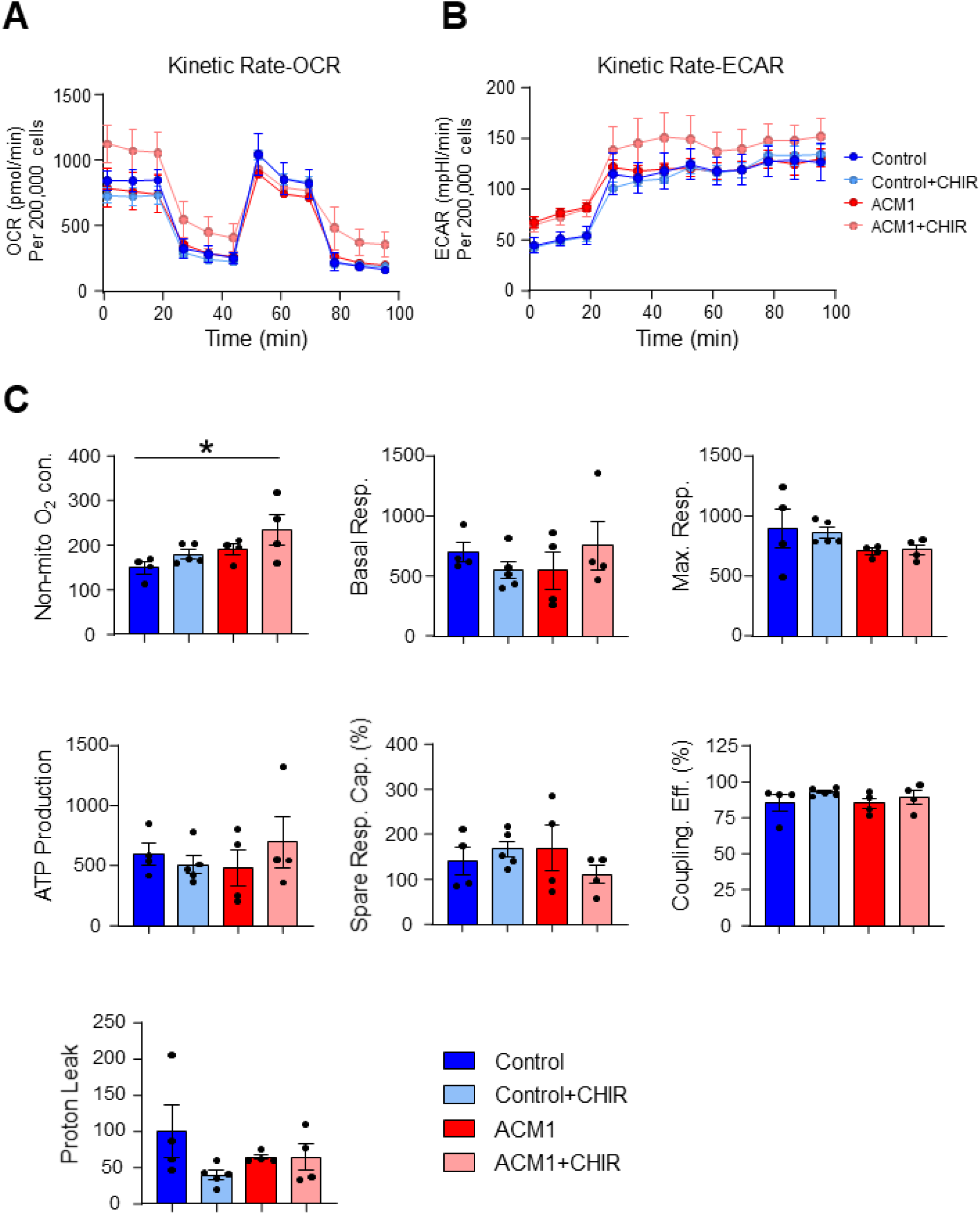
Key aspects of mitochondrial respiration are largely unchanged in ACM patient iPSC-CMs. **(A)** Real time measurements of oxygen consumption rate (OCR) and **(B)** extracellular acidification rate (ECAR) in unaffected control or ACM patient iPSC-CMs with and without GSK3 inhibition. **(C)** Analysis of various parameters of mitochondrial function in unaffected control or ACM patient iPSC-CMs with and without GSK3 inhibition. Data represent the standard error of the mean of 4 independent experiments. *, p < 0.05 compared to control.

**Figure 7.**
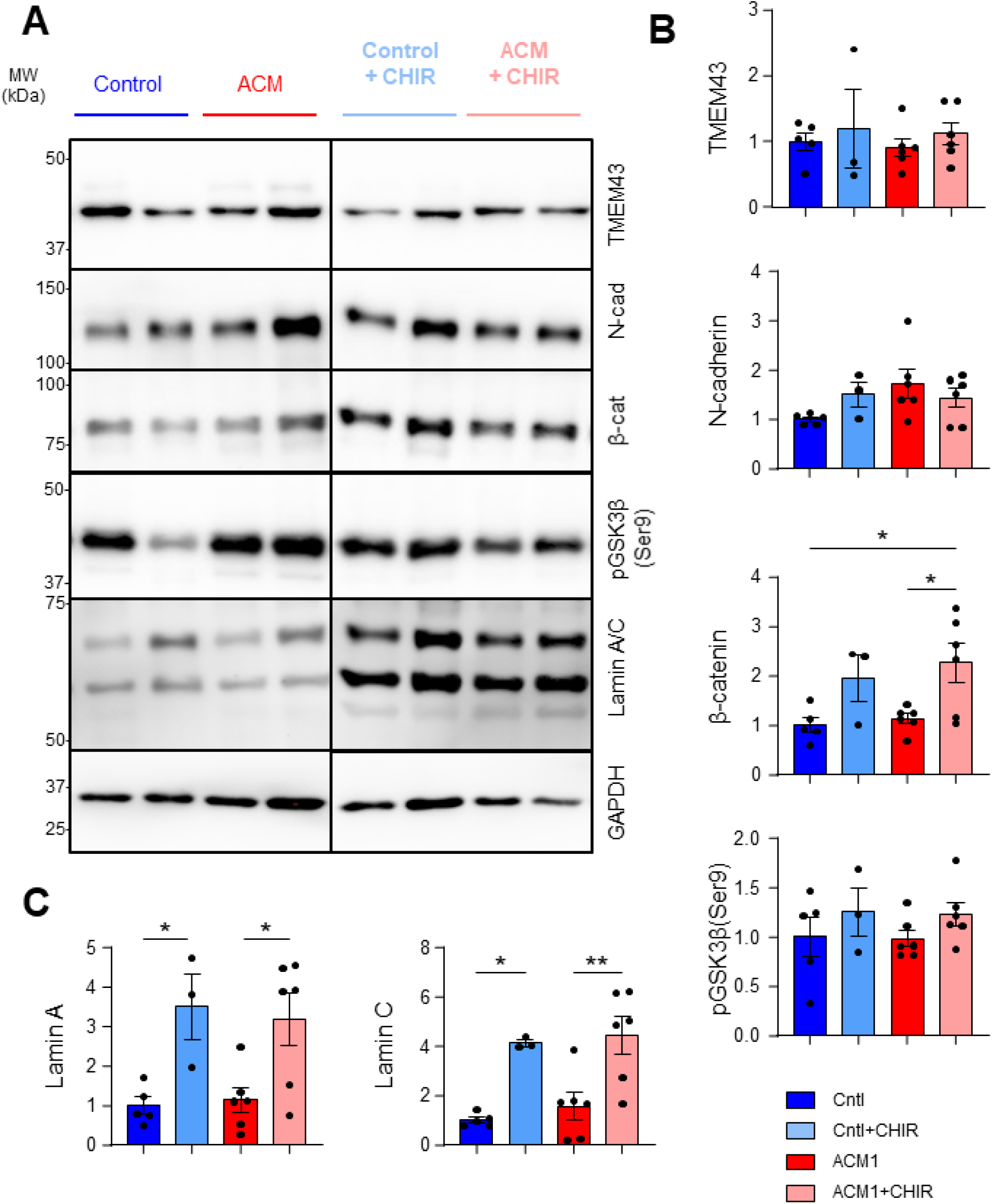
GSK3 inhibition increases β-catenin and Lamin A/C protein expression in ACM patient iPSC-CMs. **(A)** Representative Western blots and **(B, C)** densitometric analysis of various molecules in untreated and CHIR99021-treated iPSC-CMs. Data represent the standard error of the mean of 3-6 independent experiments. *, p < 0.05; **, p < 0.01 compared to untreated as determined using one-way ANOVA with Tukey’s post- hoc multiple comparisons test.

## Discussion

This is the first comprehensive characterization of iPSC-cardiomyocytes from NL ACM patients (from 27 multiplex, multigenerational families) harboring the disease-causing TMEM43 p.S358L variant^6, 11, 20^. The TMEM43 p.S358L variant is fully penetrant, in that every person who inherits the disease-causing allele will develop at least one feature of ACM over their lifespan^5, 6, 11, 12, 20, 21^. The worst outcome is SCD in young, previously asymptomatic individuals. However, clinical presentations in ACM patients are very variable, ranging from minor signs or symptoms to heart failure requiring transplant. The biggest contributor to disease variability is sex:^1^, the second is exercise^12–14^, thus the advice to children and adults with the TMEM43 p.S358L variant is to avoid high intensity sports.

While ACM caused by TMEM43 p.S358L results in both ventricular arrhythmias and cardiomyopathy, we focused our first study on the pro-arrhythmic tendencies of NL ACM patient iPSC-cardiomyocytes. iPSC-CMs from patients with different types of cardiomyopathy often display abnormal calcium handling, myofibrillar disarray, contractile dysfunction and adipogenic gene expression^22^. Our ACM patient iPSC-CMs do not appear to have obvious defects in sarcomere structure, although we did observe significantly elevated contraction frequencies and a variety of pro-arrhythmic traces in ACM patient iPSC-CMs. The most common pro-arrhythmic events we observed were alternans-like patterns, failure of the calcium to return to baseline - most likely due to EADs, and uneven event intervals (missed / irregularly spaced contractions). Future studies will examine the electrophysiology of our ACM patient iPSC-CMs to further explore the underlying causes of the observed pro-arrhythmic traces.

Although we observed no difference in phospho-GSK3β (Ser9) or phospho-AKT(Ser473) levels in ACM patient iPSC-CMs relative to control, GSK3 inhibition with CHIR99021 still conferred protection from arrhythmic contractions. GSK3β participates in many signaling pathways that regulate diverse cellular functions including glucose metabolism, protein synthesis, cell mobility, proliferation, survival, and cell fate^23, 24^ and other studies have shown beneficial effects of GSK3 inhibition for arrhythmogenic cardiomyopathy models^17, 18, 25^. Indeed, the “Targeted Therapy With Glycogen Synthase Kinase-3 Inhibition for Arrhythmogenic Cardiomyopathy (TaRGET)” clinical trial will soon begin recruiting ACM patients to evaluate the therapeutic effectiveness of the GSK3β inhibitor tideglusib (https://clinicaltrials.gov/study/NCT06174220). Our findings suggest that mitochondrial function is similar between control and ACM1 in the absence of CHIR99021 and that GSK3 inhibition with CHIR99021 does not alter key parameters of mitochondrial respiration. However, we did observe a significant increase in protein expression of β-catenin, and LaminA/C in CHIR99021-treated cells. TMEM43 has been linked to nuclear envelope integrity and thus, this may provide a link to how GSK3 inhibition confers protection in ACM patient iPSC-CMs with the TMEM43-S358L mutation^26, 27^.

The molecular function of TMEM43 protein is poorly understood and how the TMEM43 p.S358L pathogenic variant causes ACM is virtually unknown. Uncovering the fundamentals of the TMEM43 protein life cycle and localization are essential to understanding the molecular role of TMEM43 function. Future studies will continue to uncover how TMEM43 p.S358L mutation alters TMEM43 life cycle, subcellular localization, oligomerization and other protein interactions and how these parameters contribute to cardiomyocyte instability.

Efforts to date within the province of Newfoundland and Labrador have led to an average lifespan increase of 18 years for ACM patients compared to their untreated ancestors^5^. However, ACM caused by TMEM43 p.S358L mutation continues to ravage families and our lack of understanding of the molecular basis of the disease limits our ability to predict disease severity and inform patient treatment options. Future studies will examine the fibrotic or adipogenic gene expression in ACM patient samples, expand the studies to female patients, and try to understand the phenotypic variability in disease presentation and progression.

## Ethics Statement

The research performed as a part of this study were approved by the Newfoundland and Labrador Human Research Ethics Board # 2020.301 and # 2018.201.

## Author Contributions

R.J.N., W.S., and C.F. performed experiments and analysed the data. U.B. reprogrammed the iPSCs. R.J.N performed CRISPR-Cas9 gene editing. D.P. collected the dermal biopsies. K.H. recruited patients and contributed substantive ideas and advice as well as interpretation of the data. J.L.E. oversaw the project. All authors reviewed and edited the manuscript.

## Acknowledgments

We thank Dr. Dale Laird for advice about the electron micrographs. Thank you to Dr. Terry Hébert for introducing us to the Origin software for calcium imaging. We also thank Drs. Stephanie Protze and Bruno Stuyvers for their advice and suggestions. Finally, we sincerely thank the donors and their families, without whom none of this research would be possible.

## Data Availability Statement

The datasets generated during and/or analysed during the current study are available upon request

**Supplemental Videos 1 & 2.** Phase contrast movies of contracting iPSC-cardiomyocytes generated from severely affected male ACM patient and unaffected sibling control.

**Supplemental Video 3, 4 & 5.** Pseudo-coloured Fluo4 calcium fluorescence time-lapse videos of contracting iPSC-cardiomyocytes generated from 2 severely affected male ACM patients and one unaffected sibling control. Scale bar as indicated.

